# A minimal presynaptic protein machinery mediating synchronous and asynchronous exocytosis and short-term plasticity

**DOI:** 10.1101/2024.04.15.589559

**Authors:** Dipayan Bose, Manindra Bera, Chris A. Norman, Yulia Timofeeva, Kirill E. Volynski, Shyam S. Krishnakumar

## Abstract

Neurotransmitters are released from synaptic vesicles with remarkable precision in response to presynaptic Ca^2+^ influx but exhibit significant heterogeneity in exocytosis timing and efficacy based on the recent history of activity. This heterogeneity is critical for information transfer in the brain, yet its molecular basis remains poorly understood. Here, we employ a biochemically-defined fusion assay under physiologically-relevant conditions to delineate the minimal protein machinery sufficient to account for different modes of Ca^2+^-triggered vesicle fusion and short-term facilitation. We find that Synaptotagmin-1, Synaptotagmin-7, and Complexin, synergistically restrain SNARE complex assembly, thus preserving vesicles in a stably docked state at rest. Upon Ca^2+^ activation, Synaptotagmin-1 induces rapid vesicle fusion, while Synaptotagmin-7 mediates delayed fusion. Competitive binding of Synaptotagmin-1 and Synaptotagmin-7 to the same SNAREs, coupled with differential rates of Ca^2+^-triggered fusion clamp reversal, govern the kinetics of vesicular fusion. Under conditions mimicking sustained neuronal activity, the Synaptotagmin-7 fusion clamp is destabilized by the elevated basal Ca^2+^ concentration, thereby enhancing the synchronous component of fusion. These findings provide a direct demonstration that a small set of proteins is sufficient to account for how nerve terminals adapt and regulate the Ca^2+^-evoked neurotransmitter exocytosis process to support their specialized functions in the nervous system.

---

Information transfer in the brain depends on the controlled yet rapid synaptic release of neurotransmitters stored in synaptic vesicles (SVs). SV fusion and neurotransmitter release are tightly regulated by changes in the presynaptic Ca^2+^ concentration ([Ca^2+^]) and can occur in less than a millisecond after the action potential (AP) invades a presynaptic terminal^1,2^. In addition to fast, synchronous release that keeps pace with APs, many synapses also exhibit delayed asynchronous release that persists for tens to hundreds of milliseconds^1,2^. Synapses also vary in terms of how the probability of neurotransmitter release is altered by the recent history of AP firing^3,4^. The balance between synchronous and asynchronous release, and the degree of synaptic facilitation or depression, differs not only among neurons but also among synapses supplied by a single axon according to their postsynaptic targets. This heterogeneity is important in coordinating activity within neuronal networks^5–7^.

The key components of the synaptic vesicular exocytosis machinery have been identified^1,2,8^. These include the SNARE proteins that catalyze SV fusion (VAMP2 on the SV, and Syntaxin/SNAP25 on the presynaptic membrane); Ca^2+^ release sensors that couple SV fusion to Ca^2+^ signal (Synaptotagmins, Syt); and proteins that regulate SV docking and the organization of vesicular release sites (e.g., Complexin (CPX), Munc13, Munc18). Ca^2+^-evoked neurotransmitter release occurs from a readily releasable pool (RRP) of vesicles docked at the presynaptic active zone^1,2,9^. A consensus has emerged that, at an individual RRP vesicle, multiple SNARE complexes are arrested (‘clamped’) in a partially-assembled state (SNAREpins) by Synaptotagmins and CPX. Ca^2+^ activation of Synaptotagmins releases the fusion clamp allowing SNAREpins to fully assemble and drive SV fusion^10–12^. Despite this general scheme, the reasons for variations in the synchrony of exocytosis or the occurrence of short-term facilitation or depression among synapses, remain poorly understood.

Synchronous neurotransmitter release, occurring within a few milliseconds of the arrival of an AP, is triggered by a transient high local [Ca^2+^] ([Ca^2+^]_peak_ ∼ 10-100 µM) at vesicular release sites, and genetic deletion and substitution experiments have shown that it requires a fast, low-affinity Ca^2+^ sensor such as Syt1, Syt2 or Syt9^13,14^. Asynchronous neurotransmitter release can occur in response to a single AP but is particularly prominent during and following high-frequency bursts of APs. This delayed release requires a persistent elevation of presynaptic [Ca^2+^] and this [Ca^2+^]_residual_ is thought to reach a low micromolar concentration^1,15,16^. At many synapses, accumulation of [Ca^2+^]_residual_ also leads to transient facilitation of the fast synchronous release component^1,4^. The slow, high-affinity Ca^2+^ sensor Syt7, which is activated by both [Ca^2+^]_peak_ and [Ca^2+^]_residual_, has been implicated in regulating both asynchronous release and short-term facilitation^17–20^. Indeed, previous studies have shown that genetic removal of Syt1 eliminates the fast, synchronous component of evoked SV release whilst deletion of Syt7 reduces short-term synaptic facilitation and asynchronous release^13,18,19^.

An inherent limitation of genetic studies is their inability to demonstrate whether Syt1 and Syt7 alone are *sufficient* to regulate the timing and plasticity of neurotransmitter release, as the contribution of other presynaptic proteins cannot be ruled out. Furthermore, because vesicular exocytosis involves an interplay of presynaptic Ca^2+^ dynamics and Ca^2+^ sensors, a quantitative account of synchronous/asynchronous release kinetics and short-term plasticity requires precise control and measurement of [Ca^2+^]. This is difficult to achieve in intact synapses owing to the small size of the active zone and the high speed of Ca^2+^ diffusion and buffering^21,22^.

Hence, we sought to determine whether Syt1 and Syt7 (along with SNAREs and CPX) are sufficient to determine the timing and activity-dependent plasticity of SV release. Additionally, we aimed to uncover the underlying molecular mechanisms governing the cooperative action of Syt1 and Syt7. To achieve this, we took a reductionistic approach of combining an *in vitro* reconstituted fusion assay^23–26^ with quantitative computational modeling^12^. Specifically, we utilized a biochemically-defined high-throughput assay based on a suspended lipid membrane platform that uses fluorescence microscopy to track the docking, clamping (equivalent to the delay from docking to spontaneous fusion), and Ca^2+^-triggered fusion of individual vesicles at tens of milliseconds precision. Critically, this setup allowed precise control over the identity and density of the included proteins, as well as the [Ca^2+^] signal^23,25–27^.

Recently, using this *in vitro* experimental setup, we demonstrated that under physiologically relevant conditions, Syt1 and CPX are sufficient to produce clamped (RRP-like) vesicles, and these stably docked vesicles can be triggered to fuse rapidly by Ca^2+^ addition^27^. Building on this advance, we designed the reconstitution conditions to investigate the cooperative action of Syt1 and Syt7 as follows: in all experiments, we used small unilamellar vesicles containing VAMP2 and Syt1 and included CPX in the solution (Figure 1A, Supplementary Figure 1). We reconstituted pre-formed t-SNAREs (a 1:1 complex of Syntaxin1 and SNAP-25) and Syt7 (when warranted) in the suspended lipid membrane (Figure 1A, Supplementary Figure 1). In all cases, we monitored large ensembles of vesicles (∼150 - 200 vesicles) and used fluorescently labeled lipid (2% ATTO647N-PE), introduced in the vesicles to track the docking and fate of individual vesicles (Figure 1A and Materials and Methods). To trigger the fusion of docked vesicles, we chose a final [Ca^2+^] of 100 µM. This concentration aligns with the [Ca^2+^]_peak_ observed at presynaptic vesicular release sites^21^ and is sufficient to saturate both Syt1 and Syt7^10^, therefore mitigating possible variability stemming from differential activation of Syt1 and Syt7.

**Figure 1.**
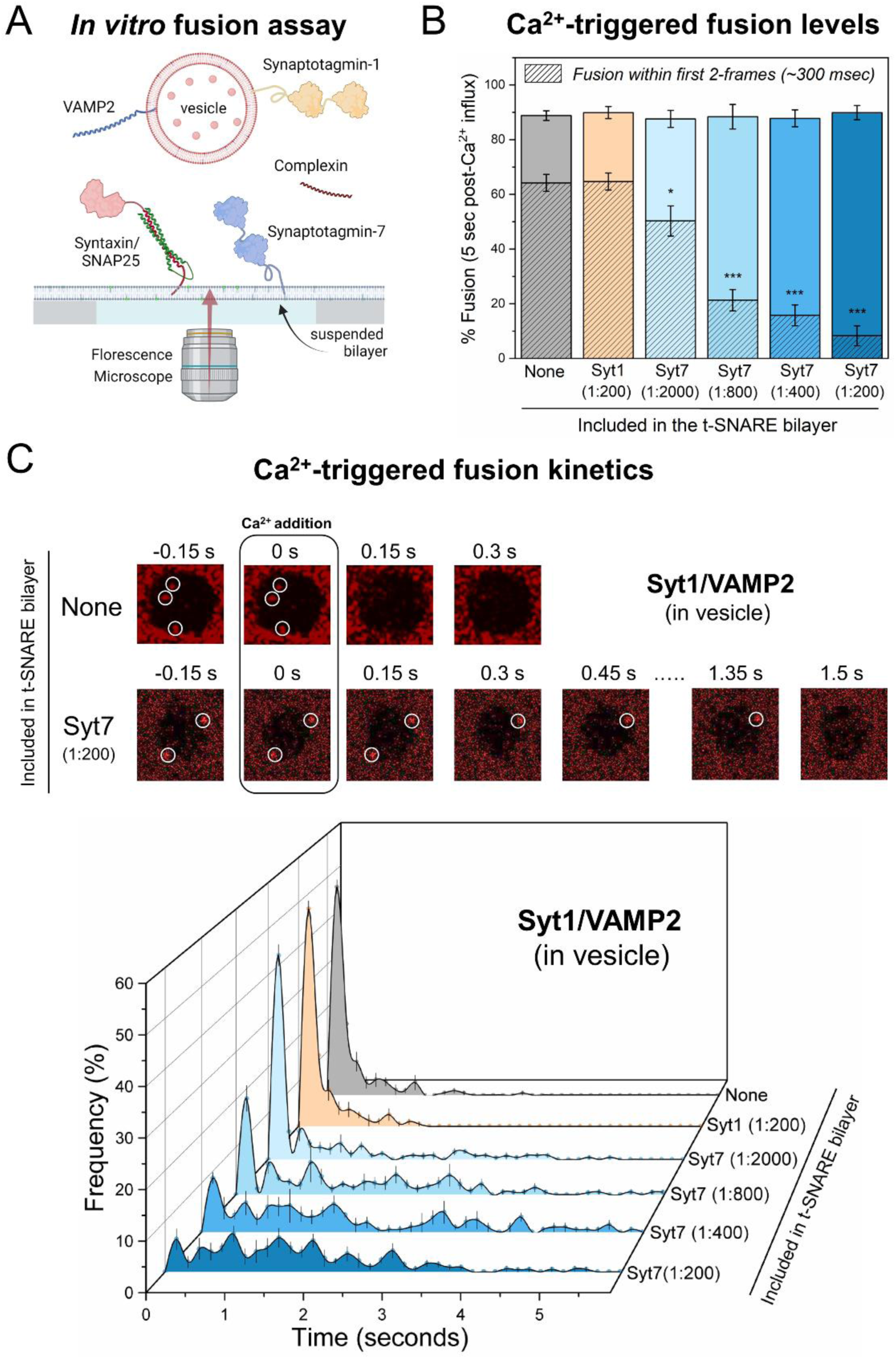
Impact of Syt7 on Ca^2+^-evoked fusion of Syt1/VAMP2 vesicles. (A) In a typical *in vitro* fusion experiment, vesicles containing VAMP2 (∼70 copies) and Syt1 (∼20 copies) were added to a suspended bilayer membrane (formed on a silicon substrate with 5 μm holes) reconstituted with Syntaxin/SNAP25 (1:400 protein-to-lipid ratio) ± Syt7. The fate of each vesicle before and after the addition of 100 µM Ca^2+^ was monitored by a confocal microscope using a fluorescent marker included in the vesicle. (B) Syt7 (included in the t-SNARE containing bilayer) had no impact on the fusion competence of docked Syt1/VAMP2 vesicles, with ∼85% fusing within 5 seconds after the arrival of 100 μM Ca^2+^ signal at or near the docked vesicles. The hatched bar represents the percent fusion occurring with 2 frames (∼300 ms) following Ca^2+^ arrival. (C) Syt7 altered the Ca^2+^-triggered fusion kinetics of docked Syt1/VAMP2 vesicles. Top, Representative time-lapse image of Ca^2+^-evoked fusion of docked vesicles shows that without Syt7, the vesicles fuse rapidly and synchronously following Ca^2+^ addition. The inclusion of Syt7 (1:200 protein-to-lipid ratio) introduces variable delays in Ca^2+^-evoked fusion kinetics. Bottom, quantitative analysis of Ca^2+^-evoked fusion of Syt1/VAMP2 vesicles shows that Syt7 introduces a concentration-dependent delay in the Ca^2+^-evoked fusion kinetics, resulting in a significant reduction in the proportion of vesicles fusing within the first 2 frames (∼300 ms) following Ca^2+^ arrival at time t = 0 seconds. Data (mean ± standard deviation) are from 5 independent experiments for each condition (∼ 40 – 50 vesicles per experiment). *p<0.05, *** p<0.001 using the Student’s t-test comparison to the condition without Syt7 in the bilayer.

## Syt7 delays the fusion of Syt1-containing vesicles in a concentration-dependent manner

In the absence of Syt7, the majority (∼95%) of the Syt1/VAMP2 containing vesicles that docked to the t-SNARE bilayers were ‘immobile’ and remained unfused during an initial 10 min observation window (Supplementary Figure 2). Addition of Ca^2+^ (100 µM) triggered the fusion of ∼90% of the stably clamped vesicles within 5 seconds as measured by lipid mixing (Figure 1B). Notably, a significant portion of fusion (∼70%) occurred within 2 frames (∼300 ms) following the arrival of the Ca^2+^ signal at or near the docked vesicles (Figure 1B, C).

Inclusion of Syt7 in the t-SNARE-containing bilayer (at concentrations ranging from 1:2000 to 1:200 protein-to-lipid ratio) had no discernable effect on the number or the fate of the docked Syt1/VAMP2 vesicles (Supplementary Figure 2). Hence, the vast majority (∼90%) of the vesicles remained stably docked in an immobile clamped state (Supplementary Figure 2). Likewise, Syt7 did not impact the Ca^2+^-induced fusion competence as ∼90% of the docked vesicles fused within 5 seconds following the addition of Ca^2+^ (100 µM) (Figure 1B). However, we observed significant delays in the kinetics of Ca^2+^-triggered fusion, and these delays correlated with the amount of Syt7 included in the bilayer (Figure 1C). The proportion of ‘coupled release’ i.e. vesicles undergoing fusion within 2 frames (∼300 ms) following the arrival of Ca^2+^ signal progressively declined from approximately 70% to 10%, as the concentration of Syt7 in the bilayer was increased from 1:2000 to 1:200 (Figure 1B, C). Noteworthy, this impact was exclusive to Syt7, as inclusion of Syt1 (instead of Syt7) in the suspended bilayer, even at a protein-to-lipid ratio of 1:200, did not alter the likelihood or kinetics of Ca^2+^-triggered fusion of Syt1/VAMP2 vesicles (Figure 1B, C). Taken together, these data indicate that Syt7 influences the kinetics of Ca^2+^-triggered fusion for Syt1/VAMP2 vesicles in a concentration-dependent manner, while not affecting the fusion competence of docked vesicles.

## Syt1 and Syt7 cooperate to establish the fusion clamp at rest

The clamping efficiency of docked vesicles was similar in the absence or presence of Syt7, with approximately 90% of docked vesicles stably clamped under both conditions (Supplementary Figure 2). Therefore, it remained unclear whether Syt7 also contributes to the fusion clamp in these conditions. Hence, we developed reconstitution conditions specifically tailored to investigate the role of Syt7 in the fusion clamp. Simple removal of Syt1 and/or CPX from the reaction mixture was not feasible, as omitting CPX potentiated spontaneous fusion, while leaving out Syt1 significantly reduced the number of docked vesicles, precluding any meaningful analysis^23,27^.

In previous work, we demonstrated that under low VAMP2 copy number conditions (i.e. vesicles containing ∼13 copies of VAMP2 and ∼22 copies of Syt1), Syt1 alone could produce stably clamped Ca^2+^-sensitive vesicles^24,27^. Furthermore, disrupting the interaction between Syt1 and t-SNARE through mutations in the Syt1 C2B domain (R281A, E295A, Y338W, R398A, R399A; referred to as Syt1^Q^) abolished the Syt1 fusion clamp^27^. Given this background, we investigated Syt7’s impact on the fusion clamp by utilizing the Syt1^Q^ mutant in the CPX-free, low VAMP2 condition (Figure 2).

**Figure 2.**
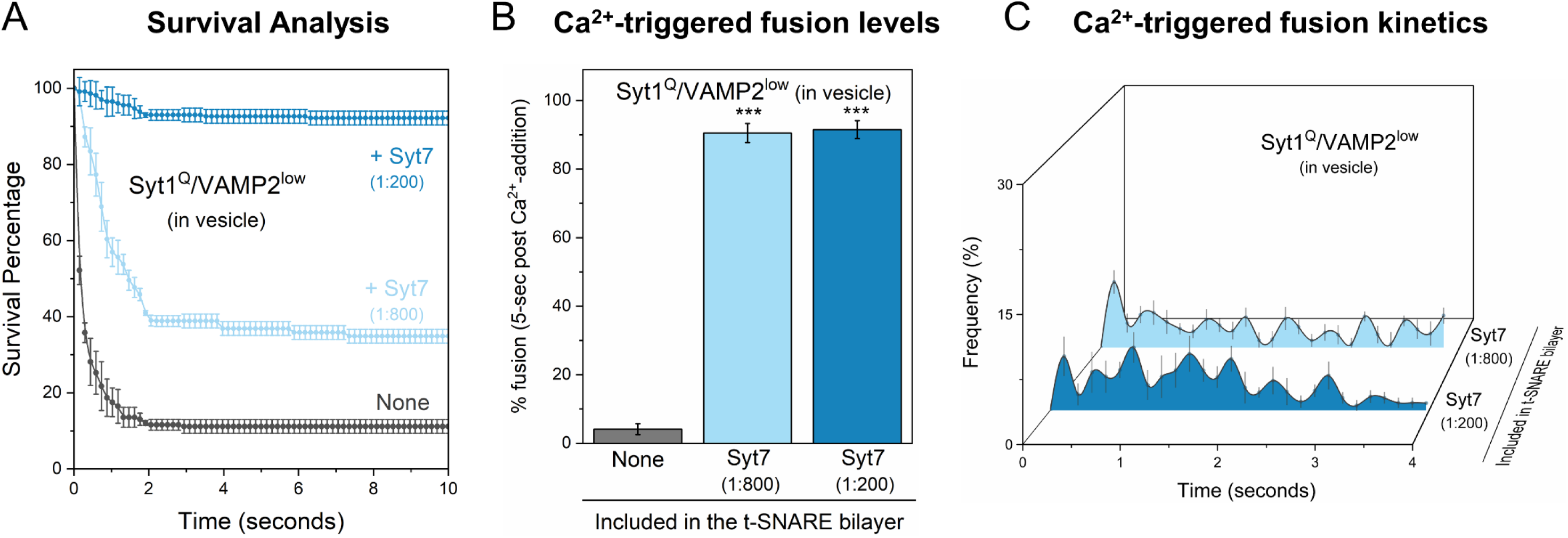
Contribution of Syt7 to the establishment of the fusion clamp. The involvement of Syt7 in the fusion clamp was evaluated using vesicles containing low-copy VAMP2 (∼15 copies) and a non-clamping Syt1 mutant, Syt1^Q^ (carrying R281A,E295A,Y338W,R398A,R399A mutations that disrupt the Syt1-SNARE primary interface) in the absence of CPX. (A) The time between docking and spontaneous fusion was measured for each docked vesicle and the ‘docking-to-fusion’ latency time was cumulatively expressed as the survival percentage. This ‘survival analysis’ provided the measure of the strength of the fusion clamp. In the absence of Syt7 (grey), the majority of the docked VAMP2^low^/Syt1^Q^ vesicles proceed to fuse spontaneously with a half-time of ∼1 sec. The inclusion of Syt7 in the bilayer resulted in stably docked vesicles in an immobile state, with clamping efficiency correlating with the amount of Syt7 included. Approximately 40% of vesicles were clamped under low Syt7 concentration (1:800, light blue) and this increased to ∼90% under high Syt7 concentration (1:200, dark blue). (B, C) Syt7 clamped VAMP2^low^/Syt1^Q^ vesicles remained fusion competent and could be triggered to fuse by the addition of Ca^2+^ (100 µM) and the observed fusion was desynchronized to the Ca^2+^ signal. In the absence of Syt7, a very small percent of the docked VAMP2^low^/Syt1^Q^ vesicles underwent fusion which precluded meaningful kinetic analysis. Data (mean ± standard deviation) are from 4 independent experiments for each condition (∼ 40 – 50 vesicles per experiment).*** p<0.001 using the Student’s t-test comparison to the condition without Syt7 in the bilayer.

As anticipated, in the absence of Syt7, the majority (>90%) of docked Syt1^Q^ vesicles fused spontaneously. Inclusion of Syt7 in the bilayer restored the clamp on Syt1^Q^ vesicles in a dose-dependent manner (Figure 2A). Approximately 40% and 90% of vesicles remained stably docked in an immobile state with Syt7 included at 1:800 and 1:200 (protein-to-lipid ratio) respectively (Figure 2A). Furthermore, these stably docked vesicles could be triggered to fuse by the addition of 100 µM Ca^2+^ (Figure 2B), but the fusion kinetics were desynchronized from the Ca^2+^ signal, with a temporally distributed vesicle fusion pattern (Figure 2C). This indicates that Syt7 can independently establish a Ca^2+^-sensitive fusion clamp. It further suggests that, under physiologically relevant conditions, Syt7 acts in concert with Syt1 and CPX to arrest SNARE assembly and produce stably, docked vesicles at rest.

## Syt1 and Syt7 regulate Ca^2+^-evoked fusion via competitive binding to the SNARE complex

Next, we investigated the impact of Ca^2+^-binding-deficient Syt1 and Syt7 mutants to understand the mechanisms behind their synergistic action (Figure 3). Specifically, we employed Syt1 with D309A, D363A, D365A mutations in the C2B domain (Syt1^DA^), and Syt7 with D225A, D227A, D233A, D357A, D359A mutations in the C2AB domains (Syt7^DA^). The introduction of Ca^2+^-insensitive Syt7^DA^ in the bilayer resulted in a concentration-dependent reduction in Ca^2+^-evoked fusion of vesicles containing Syt1^WT^/VAMP2, with ∼30% decrease at a low (1:800 protein-to-lipid) and ∼70% reduction at high (1:200) Syt7^DA^ concentrations (Figure 3A). However, Syt7^DA^ had no discernable effect on the fusion kinetics, with the majority of vesicles fusing within the first 2 frames following Ca^2+^ arrival (Figure 3A).

**Figure 3.**
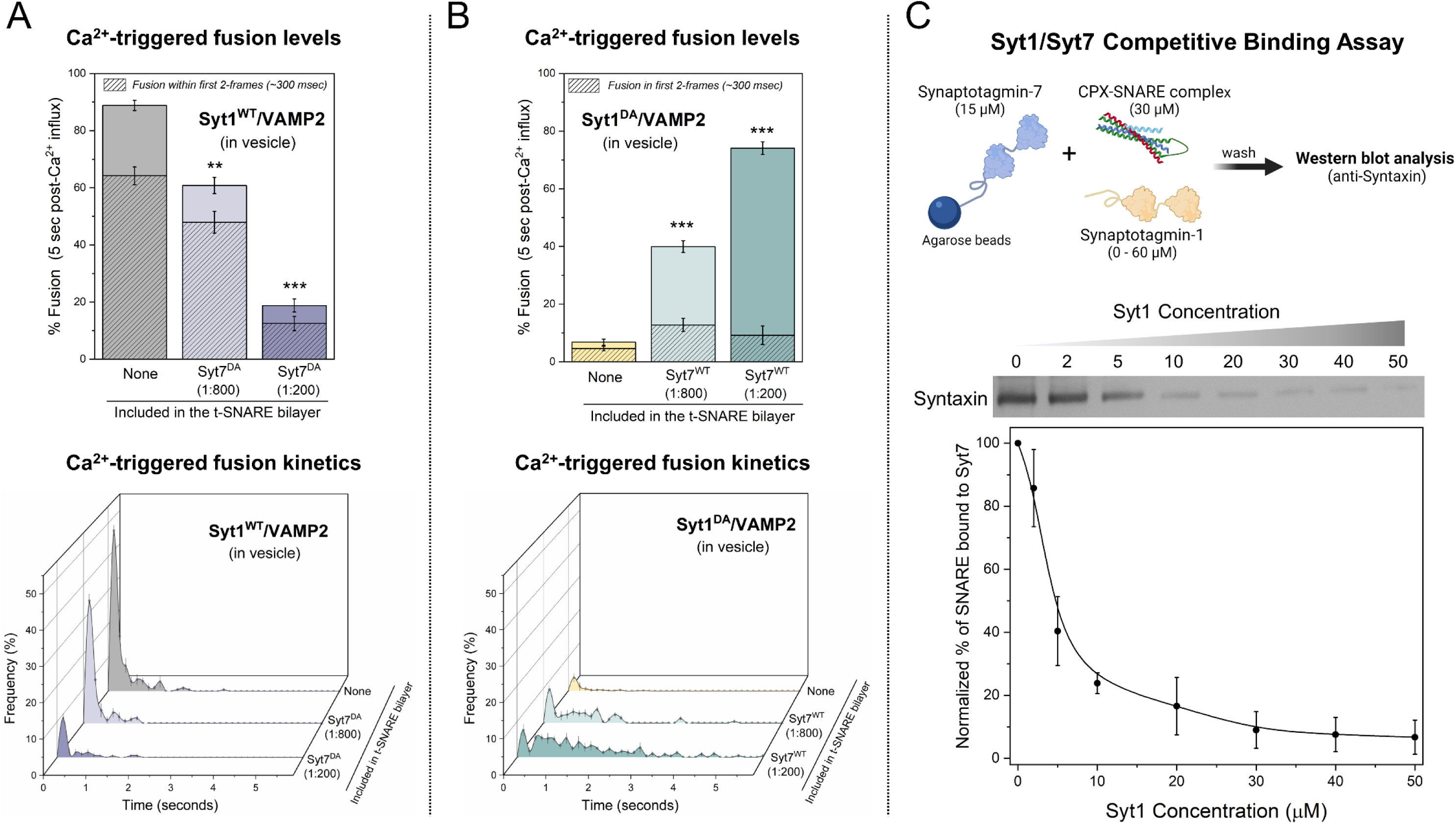
Syt1 and Syt7 synergistically regulate Ca^2+^-evoked fusion via competitive binding to the same SNARE complex. (A) The inclusion of the Ca^2+^ binding deficient Syt7 mutant (Syt7^DA^) in the bilayer inhibited Ca^2+^ (100 µM) evoked fusion of Syt1^WT^/VAMP2 vesicles in a dose-dependent manner, without altering the overall fusion kinetics. (B) Syt7^WT^ from the bilayer rescued the Ca^2+^-evoked fusion of Syt1^DA^ /VAMP2 vesicles but the fusion events were desynchronized to the Ca^2+^ signal. (C) Quantitative pull-down and Western-blot analysis with Syt7 as ‘bait’ and CPX-SNARE complex as ‘prey’ reveal that Syt1 disrupts Syt7-SNARE interaction in a concentration-dependent manner. Data (mean ± standard deviation) are from 5 independent experiments for each condition (∼ 40 – 50 vesicles per experiment) in A and B and from 3 independent experiments in C. **p<0.01, *** p<0.001 using the Student’s t-test comparison to the condition without Syt7 in the bilayer.

As expected, the disruption of Ca^2+^ binding to Syt1 (Syt1^DA^) eliminated Ca^2+^-triggered vesicular fusion (∼7%). However, the inclusion of Syt7^WT^ into the bilayer restored Ca^2+^-evoked fusion to levels corresponding with the concentration of Syt7^WT^ in the bilayer (∼40% and ∼75% with 1:800 and 1:200 Syt7^WT^ respectively) (Figure 3B). Notably, the observed fusion was desynchronized from the Ca^2+^ signal (Figure 3B). Taken together, these data suggest that Syt1 and Syt7 act on the same vesicles, likely targeting the same SNARE complexes, and their cooperative action in regulating Ca^2+^-evoked fusion stems from a competitive binding of Syt1 and Syt7 to the same SNARE complex. Furthermore, these results demonstrate that Syt1 acts as a ‘fast’ Ca^2+^-sensor to trigger rapid Ca^2+^-evoked vesicle fusion, whereas Syt7 functions as a ‘slow’ Ca^2+^-sensor that mediates release over longer time intervals.

Subsequently, we employed a quantitative pull-down assay to directly test the competitive interaction between Syt1 and Syt7 with the same SNARE complex. While the binding of Syt1 to Syntaxin/SNAP25 is well-documented^28–30^, the Syt7-SNARE interaction remains poorly understood. Hence, we initially conducted a pull-down experiment using Syt7 immobilized on agarose beads as ‘bait’ and pre-formed CPX-SNARE complex at varying concentrations as the ‘prey’. Western-blot analysis confirmed direct molecular interaction between the CPX-SNARE complex and Syt7, revealing a saturable dose-response curve with an estimated apparent affinity (K_d_) ∼ 20 µM (Supplementary Figure 3). We then examined the binding of 30 µM CPX-SNARE complex to Syt7-coated beads in the presence of varying concentrations (ranging from 1 μM - 50 µM) of Syt1. The inclusion of Syt1 disrupted the Syt7-SNARE interaction, resulting in near complete abrogation of binding at Syt1 concentrations ≥ 30 µM (Figure 3C). This analysis directly demonstrates the competitive nature of the binding between Syt1 and Syt7 to the SNARE complex. In summary, our data argue that the kinetics of Ca^2+^-triggered fusion are governed by the number of Syt1 or Syt7 associated SNAREpins, which is in turn determined by the relative abundance of these two proteins.

## Differential clamp removal rates of Syt1 and Syt7 shape Ca^2+^ triggered fusion kinetics

How do Syt1 and Syt7 shape the kinetics of vesicular fusion? Our data support the hypothesis that the cooperative action of Syt1 and Syt7 in regulating vesicular release can be explained by a ‘release of inhibition’ model^10–12^. According to this model, Syt1 and Syt7 along with CPX bind to SNAREpins at docked SVs and clamp vesicular fusion at rest. Ca^2+^ activation of Syt1 and Syt7 leads to the release of the fusion clamp. Thereby, the rate of the Ca^2+^-triggered removal of the fusion clamp determines the overall efficacy and kinetics of SV fusion (Figure 4A). The model further posits that the differential Ca^2+^/membrane binding properties of Syt1 and Syt7, along with the relative numbers of Syt1 or Syt7 bound SNAREs on a given vesicle, fine-tune the release properties in response to Ca^2+^ signals.

**Figure 4.**
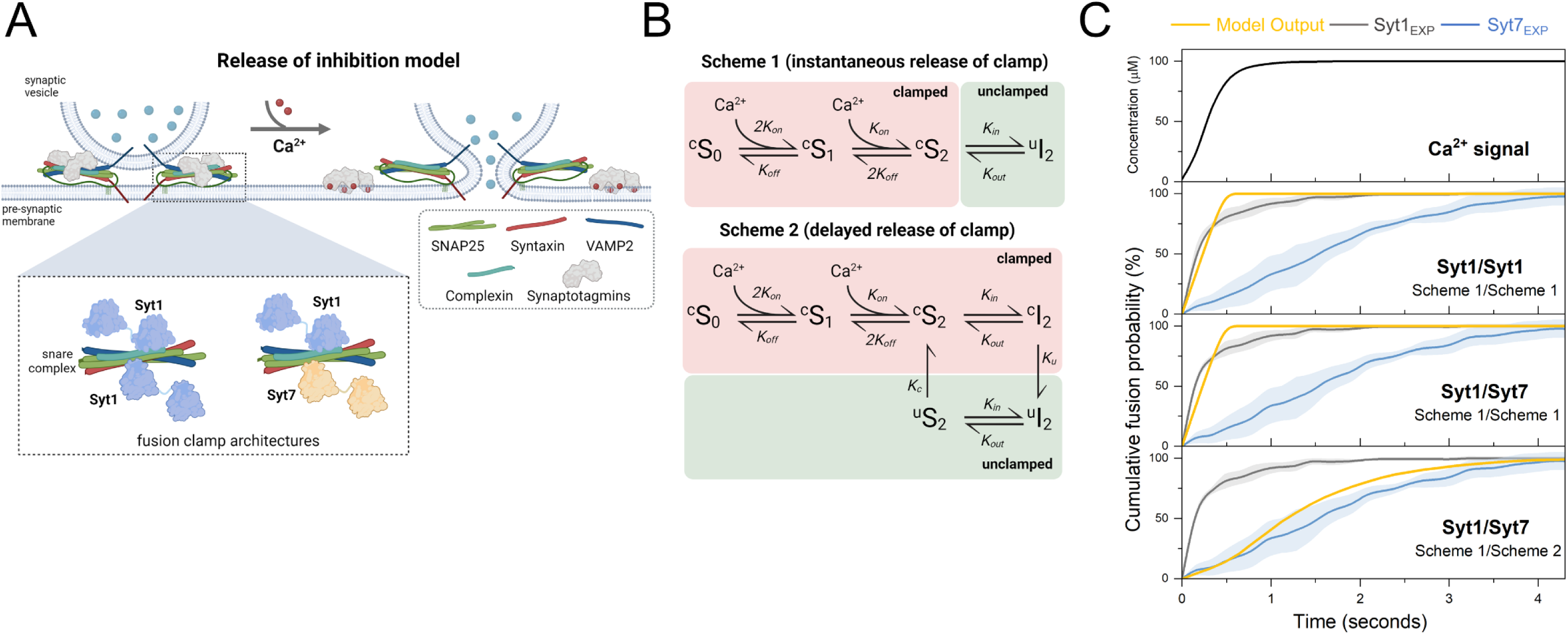
Computational model of synergistic activation of vesicular fusion by Syt1 and Syt7. (A) Schematic illustration of the release of inhibition model. At rest, fusion of vesicles is inhibited (‘clamped’) by binding of Syt1 and Syt7 along with CPX to partially assembled SNAREpins. Upon Ca^2+^ binding, the C2 domains of Syt1 and Syt7 insert into the membrane, leading to the removal of the fusion clamp. This allows the complete zippering of the SNARE complexes, resulting in vesicular fusion. Inset shows that two clamp architectures are considered in the default model: dual Syt1/Syt1 or dual Syt1/Syt7 clamp (see Supplementary Figure 5 for additional clamp architectures tested). (B) Kinetic reaction schemes describing Ca^2+^-triggered release of the fusion clamp. Each modeled C2 domain sequentially binds two Ca^2+^ ions which triggers the insertion of its aliphatic loop into the membrane. Scheme 1 assumes that membrane insertion results in the instantaneous removal of the Synaptotagmin fusion clamp, while Scheme 2 assumes a delay between membrane insertion and the removal of the clamp. (C) The time course of vesicular fusion (Model Output) simulated in response to the experimentally constrained Ca^2+^ signal (Supplementary Figure 4) for models with different clamp architecture and kinetics of clamp reversal. Experimental data (from Figure 1) for the Ca^2+^-triggered fusion of Syt1 containing vesicles in the absence (Syt1_EXP_) or the presence of saturating levels of Syt7 (Syt7_EXP_) are plotted for comparison. The model suggests that experimentally observed fusion kinetics can be explained by the mechanism with differential rates of fusion clamp removal for Syt1 (instantaneous) and Syt7 (delayed). For each modeled condition a minimum of 1000 stochastic simulations were performed to calculate the average response.

To investigate whether the differences in Ca^2+^/membrane binding properties of Syt1 and Syt7 could explain our results, we utilized the computational implementation of the release of inhibition model ^12^. This modeling framework enables us to simulate SV fusion in response to varying Ca^2+^ signals for different synaptotagmin fusion clamp architectures. Drawing on structural studies, within the default model, we assumed that each vesicle contains six SNARE complexes^31^ and each SNARE complex can bind two Syt1 molecules^28^. Additionally, we postulated that Syt7 might compete with Syt1 for one of these binding sites. Consequently, we considered two limiting cases for the fusion clamp’s architecture: either Syt1/Syt1 or Syt1/Syt7 (Figure 4A). These scenarios correspond to experimental conditions without Syt7 or with a saturating level of Syt7 in the lipid bilayer respectively. As in our previous work, we assumed that Ca^2+^ binding and membrane loop insertion of the C2B domain of Syt1 or C2A domain leads to the instantaneous removal of the fusion clamp (Figure 4B, Scheme 1). The release of the clamp enables the full zippering of freed SNAREs, and each SNARE complex independently contributes ∼4.5 kBT of energy to lower the fusion barrier, thereby catalyzing SV fusion.

As a model input, we incorporated experimentally estimated changes in [Ca^2+^] at the lipid bilayer, corresponding to a ramped increase of [Ca^2+^] from 0 to 90 µM within 0.5 seconds (Supplementary Figure 4). In the absence of Syt7 (Syt1/Syt1 clamp), the standard model closely reproduced the kinetics of vesicular fusion in response to the addition of Ca^2+^ (Figure 4C). Indeed, the time course of Syt1-mediated vesicular fusion closely follows the kinetics of the [Ca^2+^] signal (Figure 4C). This suggests that Ca^2+^ diffusion is likely the rate-limiting step governing vesicular fusion kinetics under our experimental conditions. Notably, the same computational implementation of the ‘release of inhibition’ model reproduced the millisecond kinetics of vesicular fusion in response to fast [Ca^2+^] transients that occur in live synapses^12^. This indicates that the reconstituted fusion assays mirror the functionality of these proteins within living synapses, with comparable operational efficacy.

However, this model failed to replicate the slower vesicular fusion kinetics observed when Syt7 was included (Syt1/Syt7 clamp). Indeed, under our experimental conditions, Syt1 and Syt7 exhibit comparable activation patterns, as their rates of Ca^2+^ binding and membrane association are similar under saturating [Ca^2+^]. This suggests that the Ca^2+^-triggered membrane insertion of Syt7 is not the rate-limiting step in the removal of the Syt7 fusion clamp. Consequently, we adapted the model to include a delay between the Ca^2+^-triggered membrane insertion of the Syt7 C2A domain and the removal of the fusion clamp (Figure 4B, Scheme 2). This modification allowed us to reconcile the model with the experimental data under Syt1/Syt7 clamp conditions (Figure 4C).

Given the ongoing debate surrounding the exact number of SNARE complexes on an RRP vesicle^31,32^, as well the architecture of the Synaptotagmin fusion clamp^27–29^, we explored alternative fusion clamp configurations. Specifically, we varied the number of SNAREpins in the vesicles from six to twelve and examined scenarios where a single Synaptotagmin molecule - either Syt1 or Syt7 - could bind to and clamp an individual SNAREpin (Supplementary Figure 5). In all instances, the model outputs closely matched those from the default dual-clamp model with six SNAREpins (Supplementary Figure 5). Taken together, our data argue that the mechanisms of clamp removal are distinct for Syt1 and Syt7 and that the differing rates of clamp removal – rapid for Syt1 and slower for Syt7 – are key factors determining the Ca^2+^ triggered vesicular fusion kinetics.

Both Syt1 and Syt7 are expected to bind and clamp the SNARE complexes via their C2B domains^28,30^. However, the critical distinction in their roles in the regulation of SV fusion arises from the Ca^2+^ activation of the Syt1 C2B domain compared to the Syt7 C2A domain^19^. The fast removal of the Syt1 clamp may be attributed to the rapid dissociation of Syt1 from the SNARE complex upon Ca^2+^-triggered membrane insertion of its C2B domain, as demonstrated in biochemical and structural studies^29,30^. In contrast, for Syt7 the decoupling of SNARE binding (C2B domain) and Ca^2+^ activation (C2A domain) may contribute to the slower disassembly of the Syt7 fusion clamp. Additional research into the Syt7/SNARE interaction in needed to understand the molecular mechanism of Ca^2+^-triggered removal of the Syt7 fusion clamp.

## Syt7 enhances Ca^2+^ synchronized fusion under elevated basal Ca^2+^ conditions

In addition to modulating fusion kinetics, Syt7 has also been implicated in the facilitation of synchronous neurotransmitter release during neuronal activity^17,18^. This short-term plasticity of vesicular release has been linked to the accumulation of [Ca^2+^]_residual_ in the presynaptic terminal due to sustained neuronal activity^1,15,16^. Hence, we investigated the effect of elevated basal [Ca^2+^]_basal_ on Ca^2+^-triggered release properties of Syt1/VAMP2 vesicles in the absence and presence of Syt7.

The inclusion of 0.5 μM [Ca^2+^]_basal_ during the vesicle docking phase had little to no effect on vesicle docking or clamping (i.e., spontaneous fusion) of vesicles across all conditions tested (Supplementary Figure 6). It also did not affect vesicle fusion triggered by 100 μM Ca^2+^ under control (no Syt7 in the bilayer) conditions (Figure 5A). However, when Syt7 was included in the bilayer (at 1:200 protein-to-lipid ratio), it enhanced the fast component of Ca^2+^-evoked release (within the first 300 ms after the arrival of 100 µM Ca^2+^ signal), increasing it from ∼8% to ∼35%, without changing the overall level of fusion over the 5-second interval (Figure 5A). Likewise, the computational model incorporating a delay in the removal of the Syt7 fusion clamp (Scheme 2) also replicated the synchronization of vesicular fusion when Syt7 was pre-activated with low micromolar [Ca^2+^] (Figure 5B). Importantly, the degree of synchronization correlated with the concentration of Ca^2+^ utilized for pre-activation (Figure 5B). Together, these data show that when pre-activated by low micromolar [Ca^2+^]_basal_, Syt7 enhances Ca^2+^-synchronized vesicle fusion. While the precise mechanism of Syt7-mediated facilitation is not fully understood, it is hypothesized that activation of Syt7 could enhance the release by two different mechanisms: (i) by increasing the probability of RRP vesicles and/or (ii) by enhancing the activity-dependent docking of SVs^4,33^. Our *in vitro* reconstitution experiments and modeling demonstrate that, when pre-activated, Syt7 facilitates the synchronous release of pre-docked vesicles.

**Figure 5.**
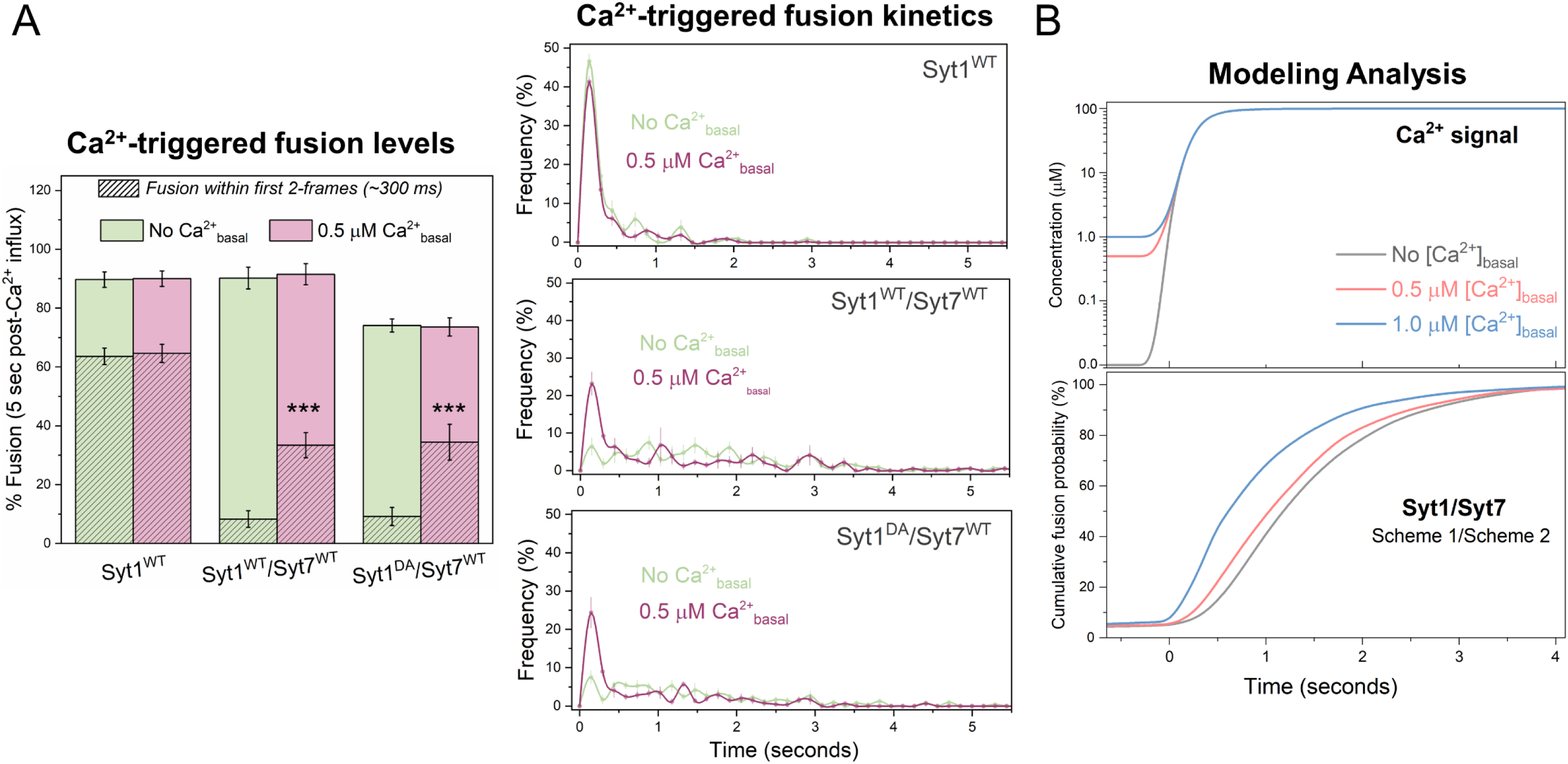
Syt7 enhances Ca^2+^-synchronized fusion under elevated [Ca^2+^]_basal_ conditions. (A) Comparison of the Ca^2+^ (100 µM) evoked fusion characteristics without (green) or with (pink) 0.5 μM [Ca^2+^]_basal_ included during vesicle docking reveals that when pre-activated, Syt7 (included in the bilayer at 1:200 protein-to-lipid ratio) increases the proportion of Ca^2+^-coupled release of Syt1-containing vesicles (Syt1^WT^/Syt7^WT^) without changing the overall fusion levels. This enhancement was not observed with Syt1^WT^ alone. Notably, a similar degree of facilitation of Ca^2+^-synchronized release was observed with vesicles containing Ca^2+^-binding deficient Syt1^DA^ (Syt1^DA^/Syt7^WT^). Data (mean ± standard deviation) are from 4 independent experiments for each condition (∼ 40 – 50 vesicles per experiment). (B) The dual Syt1/Syt7 clamp model, incorporating the delayed release of the clamp for Syt7, reproduces the experimentally observed facilitation of synchronous release following pre-activation with low micromolar [Ca^2+^]. The extent of facilitation correlated with the level of pre-activating [Ca^2+^] used. For each modeled condition a minimum of 1000 stochastic simulations were performed to compute the average response. *** p<0.001 using the Student’s t-test comparison to the condition without Syt7 in the bilayer.

Interestingly, disrupting Ca^2+^-binding to Syt1 (Syt1^DA^) did not abolish the Syt7-dependent synchronization of vesicular release with elevated [Ca^2+^]_basal_ (Figure 5A). Indeed, we observed a similar proportion of Ca^2+^-synchronized release between Syt1^WT^/Syt7 and Syt1^DA^/Syt7 conditions (Figure 5A). This indicates that when primed by elevated [Ca^2+^]_basal_, Syt7 is capable of independently mediating fast, Ca^2+^-synchronized release. Our previous computational modeling study provided insights into the possible molecular mechanisms underlying Syt7-mediated facilitation^12^. It revealed that the rate of vesicle fusion is dictated by the time taken for three SNAREpins to be released from the fusion clamp on a specific vesicle. Due to its high Ca^2+^/membrane affinity, Syt7 is partially activated at low micromolar [Ca^2+^], weakening the fusion clamp. This in turn, accelerates the liberation of three SNAREpins upon the arrival of [Ca^2+^] signal, thereby enhancing the synchronous release component. Consistent with this hypothesis, we observed a comparable level of facilitation when [Ca^2+^]_basal_ levels were elevated either during or after the vesicles reached the immobile clamped state (Supplementary Figure 6).

## Summary

Our study presents an *in vitro* reconstitution of different modes of Ca^2+^-triggered SV fusion and short-term plasticity with minimal protein components. We demonstrate that Syt1 and Syt7, along with SNAREs and CPX, are sufficient to recapitulate synchronous and asynchronous exocytosis as well as short-term facilitation of vesicular release triggered by physiologically relevant Ca^2+^ signals. We find that Syt1 and Syt7 compete to bind the same SNARE complexes and hence the relative abundance of these Ca^2+^ sensors shape the kinetics and plasticity of vesicular fusion. Additionally, our findings suggest that the distinct actions of Syt1 and Syt7 can be attributed to the differential strength and kinetics of Ca^2+^-triggered reversal of their respective fusion clamps.

We note that, apart from the distinct Ca^2+^-release sensors, other proteins and mechanisms also play a role in regulating the timing and plasticity of neurotransmitter release. For example, the genetic deletion of Syt7 does not fully eliminate asynchronous release or short-term facilitation in some synapses^34,35^. Consequently, it is suggested that other Ca^2+^-release sensors (e.g., Syt1 and Doc2A) may also contribute to the asynchronous release component^17,36,37^. Additionally, short-term facilitation of synchronous neurotransmitter release may result from enhanced Ca^2+^ transients at release sites, driven by the saturation of Ca^2+^ buffers during repetitive activity^22,38^.

Furthermore, the strength and efficacy of neurotransmitter release is also regulated by Ca^2+^-dependent SV docking, priming, and recycling. As such, SV priming factors (e.g. RIM, Munc13) could also influence neurotransmitter release dynamics^9,39^. Nonetheless, our results highlight the central role of Syt1 and Syt7 in decoding presynaptic Ca^2+^ dynamics and translating this into complex patterns of vesicular release.

## Acknowledgments

We are grateful to Drs Dimitri Kullmann, James Rothman, and Dmitri Rusakov for reading the manuscript and providing critical feedback. This work was supported by National Institute of Health (NIH) grant NS133091 (S.S.K and K.E.V); UKRI MRC Project Grant MR/T002786/1 (Y.T. and K.E.V); UKRI BBSRC/NC3R Project Grant NC/X002233/1 (K.E.V).

## Contributions

S.S.K and K.E.V conceived the project; D.B. and M.B. carried out the *in vitro* functional analysis; C.A.N, and Y.T. contributed to the implementation of the computational model; C.N. performed all model simulations. S.S.K and K.E.V wrote the manuscript. All authors discussed the results and commented on the manuscript.

## Materials and Methods

### Proteins & Materials

In this study, we used the following clones that have been described previously^23,27^ including full-length VAMP2 (human VAMP2-His^6^, residues 1–116); full-length t-SNARE complex (mouse His^6^-SNAP25B, residues 1–206 and rat Syntaxin1A, residues 1–288); CPX (human His^6^-Complexin 1, residues 1–134); Syt1 wild-type (rat Synaptotagmin1-His^6^, residues 57-421) and mutants (D309A, D 363A, D365A; Syt1^DA^) and (R281A, E295A,Y338W,R398A,R399A, Syt1^Q^) in the same background. In addition, we created and utilized a full-length Syt7 clone, which contained rat Syt7 residues 17-403 attached to the Syt1 transmembrane domain (TMD) with a flexible 16 residue GSGS linker and a N-terminal SUMO tag (SUMO-Syt1^TMD^-Syt7). Note: We included Syt1^TMD^ to the N-terminus of full-length Syt7 to enhance the protein’s reconstitution efficiency in the membrane, while the flexible linker ensured the proper orientation of the Syt7 C2AB domain. We also generated Syt7 mutant (D225A, D227A, D233A, D357A, D359A; Syt7^DA^) in the same background. We purchased the cDNA to produce the SUMO nanobody (nanoCLAMP SMT3-A1) from Nectagen (Lawrence, KS). The lipids used in the study, including 1,2-dioleoyl-snglycero-3-phosphocholine (DOPC), 1,2-dioleoyl-sn-glycero-3-(phospho-L-serine) (DOPS), and phosphatidylinositol 4, 5-bisphosphate (PIP2) were purchased from Avanti Polar Lipids (Alabaster, AL). ATTO647N-DOPE and ATTO465-DOPE were purchased from ATTO-TEC, GmbH (Siegen, Germany) and Calcium Green conjugated to a lipophilic 24-carbon alkyl chain (Calcium Green C24) was purchased from Abcam (Cambridge, UK). All other research materials and consumables, unless specified, were purchased from Sigma-Aldrich (St Louis, MO) and Thermo Fisher Scientific (Waltham, MA)

### Protein expression and purification

All proteins were expressed and purified in a bacterial expression system as described previously^23,27^. In summary, proteins were expressed in *E. coli* BL21(DE3) cells (Novagen, Madison, WI) under 0.5 mM IPTG induction for 4 hr. Bacterial cells were pelleted and then lysed using a cell disruptor (Avestin, Ottawa, Canada) in lysis buffer containing 25 mM HEPES, 400 mM KCl, 4% Triton X-100, 10% glycerol, pH 7.4 with 0.2 mM Tris[2-carboxyethyl] phosphinehydrochloride (TCEP), and EDTA-free Complete protease inhibitor cocktail (Merck, Rahway, NJ). The resulting lysate was clarified using a 45Ti rotor (Beckman Coulter, Atlanta, GA) at 40,000 RPM for 30 min and subsequently incubated with pre-equilibrated Ni-NTA resin overnight at 4°C. The resin was washed with wash buffer containing 25 mM HEPES pH 7.4, 400 mM KCl, 0.2 mM TCEP. The wash buffer was supplemented with 1% octylglucoside (OG) for Syt1 and SNARE, and with 0.2% Triton-X-100 for Syt7. Proteins were eluted from beads using 400 mM Imidazole and their concentrations were determined using a Bradford Assay (BioRad, Hercules, CA) with BSA standard. The Syt1 and Syt7 proteins were further treated with Benzonase (Millipore Sigma, Burlington, MA) at room temperature for 1 hr with Syt1 additionally being run through ion exchange (Mono S) to remove DNA/RNA contamination. SDS-PAGE analysis was done to check the purity of the proteins and all proteins were flash-frozen in small aliquots and stored at −80°C with 10% glycerol.

### Vesicle preparation

Small unilamellar vesicles containing VAMP2 and Syt1 were prepared using rapid detergent dilution and dialysis method, followed by additional purification on discontinuous Optiprep gradient by ultracentrifugation^23,27^. To mimic synaptic vesicle lipid composition, we used 88% DOPC, 10% DOPS, and 2% ATTO647N-PE, with the protein-to-lipid input ratio of 1:100 for VAMP2 for physiological density, 1:500 for VAMP2 at low copy number, and 1:250 for Syt1. Informed by previous work^23,27^ that characterized the reconstitution efficiency and inside/outside ratio of these proteins, we estimate the vesicle contains ∼70 copies of outside facing VAMP2 and ∼20 copies of outside facing Syt1 (at physiological conditions) and ∼15 copies of VAMP2 and ∼20 copies of Syt1 (under low VAMP2 conditions). For experiments using Sulforhodamine-B loaded vesicles, VAMP2/Syt1 vesicles were prepared as described above with the addition of 30 mM Sulforhodamine-B in the dilution buffer. These vesicles were subsequently purified using a Sepharose-CL4B column to remove excess dye, instead of the Optiprep gradient method.

### Suspended lipid bilayer formation

To form the suspended lipid bilayer, we first prepared giant unilamellar vesicles (GUVs) containing t-SNARE ± Syt7 were prepared using the osmotic shock protocol as described previously^40^. To mimic the presynaptic plasma membrane, the lipid composition of the GUVs was 80% DOPC, 15% DOPS, 3% PIP2%, and 2% ATTO465-PE. The t-SNARE complex (1:1 Syntaxin/SNAP25) was included at the protein-to-lipid input ratio of 1:200 to yield a final concentration of 1:400. Incorporating the t-SNARE complex enabled us to circumvent the necessity for the SNARE-assembling chaperones Munc18 and Munc13^41^. When warranted, Syt7 was added at a protein-to-lipid input ratio of 1:50, 1:100, 1:200, and 1:1000 to yield the defined concentrations of Syt7 tested. We incorporated Syt7 in the suspended lipid membrane reflecting its predominant localization in the presynaptic membrane in vivo^42^.

Subsequently, t-SNARE (± Syt7) containing GUVs were burst on freshly plasma-cleaned Si/SiO_2_ chips decorated with a regular array of 5 µm diameter holes in HEPES buffer (25 mM HEPES, 125 mM KCl, 0.2mM TCEP, 5 mM MgCl_2_ pH 7.4). The bilayers were then extensively washed with the same HEPES buffer containing 1 mM MgCl_2_. For each experiment, the fluidity of the bilayers was verified using FRAP of the Atto-465 fluorescence (Supplementary Figure 7). As a control, we tested and confirmed that the mobility of Alexa488 labeled t-SNAREs is not affected by the inclusion of Syt7 (Supplementary Figure 7).

### Single vesicle fusion assays

The vesicle docking and fusion experiments were carried out as described previously^23,25,27^. Typically, in each experiment, approximately 100 nM lipids worth of vesicles, along with CPX (2 µM final concentration) were added using a pipette and then allowed to interact with the suspended bilayer for 5 mins. ATTO647N-PE fluorescence was used to track the fate of individual vesicles, i.e. vesicle docking, post-docking diffusion, docking-to-fusion delays, and spontaneous fusion events. Docked immobile vesicles that remained un-fused during the initial 10 min observation period were defined as ‘clamped’. Fusion was identified as a sharp, rapid decrease in fluorescence intensity, as the lipids from the vesicles diffused into the bilayer. After the initial 5-minute observation period, the excess vesicles in the chamber were removed by buffer exchange, and 100 µM CaCl_2_ was added to quantify the Ca^2+^-triggered fusion of the pre-docked vesicles. To cover large areas of the planar bilayer and simultaneously record lipid mixing in large ensembles of vesicles (∼40-50 per experiment), the movies were acquired at a speed of 147 ms per frame.

Ca^2+^ typically reached the vicinity of vesicles docked on the bilayer approximately 1-2 frames post-addition^23,27^ and this correlated with the minima of the transmittance signal (Supplementary Figure 4). For select experiments, we also included Calcium Green C24 in the bilayer to directly quantify the arrival of Ca^2+^ at the bilayer and confirmed that it matched with the transmittance signal change (Supplementary Figure 4). As Calcium-green is a high-affinity Ca^2+^ sensor (K_d_ of ∼100 nM), its fluorescence signal is typically saturated within a single frame following the arrival of Ca^2+^ at the bilayer (Supplementary Figure 4). Hence, we utilized a soluble Alexa647 dye (∼25 nM) mixed with 100 μM CaCl_2_ to track the diffusion of Ca^2+^ into the chamber. Assuming similar diffusion of Ca^2+^ and the Alexa647 dye, the changes in Alexa647 fluorescence provided a reliable indicator for estimating alterations in the [Ca^2+^] signal at or near the vesicles docked on the bilayer (Supplementary Figure 4).

All experiments were carried out at 37°C using an inverted laser scanning confocal microscope (Leica-SP5) equipped with a multi-wavelength argon laser including 488 nm, diode lasers (532 nm and 641 nm), and a long-working distance 40X water immersion objective (NA 1.1). The emission light was spectrally separated and collected by photomultiplier tubes.

### Pull-down binding analysis

To investigate the binding of Syt7 to SNAREs and assess the competitive binding of Syt1/Syt7 to the same SNARE complex, we used a pull-down analysis coupled with western-blot analysis. Briefly, we purified a SUMO-nanobody and covalently attached it to a CNBR-activated Sepharose resin. The nanobody-Sepharose resin was incubated (4 hr at 4°C) with SUMO-Syt7 protein and subjected to extensive wash with HEPES buffer (50 mM HEPES, 400 mM KCl, 0.2 mM TCEP, 0.2% Triton-X-100, pH 7.4) to form the ‘bait’. The CPX-SNARE complex was assembled and purified on the Superdex-200 column as described previously ^43,44^ in the HEPES buffer and used as the ‘prey’. For the binding experiment, ∼15 µM of Syt7-resin was incubated with pre-formed CPX-SNARE complex at varying (1 – 50 µM) concentrations overnight at 4°C with minimum agitation. The resin was washed extensively (5X) with HEPES buffer, followed by a stringent wash with HEPES buffer containing 1M KCl to eliminate unbound proteins. The resin samples were subjected to SDS-PAGE gel electrophoresis, followed by western blotting using a Syntaxin monoclonal antibody (Abcam, Cambridge, UK) to quantify the amount of SNARE bound to the Syt7-resin. We used the same protocol for the competition assay, with the following modification: 15 µM Syt7-resin was incubated with 30 µM of CPX-SNARE complex, along with 1-60 µM Syt1 included in the solution overnight at 4°C with minimum agitation.

### Computational Modelling

Vesicular fusion in response to experimentally estimated changes in [Ca^2+^] was simulated using the computational modelling framework established in our previous work^12^. [Ca^2+^] stimulation profile was approximated based on the diffusion kinetics of Alexa 647 (Supplementary Figure 4). Each RRP vesicle was associated with either six or twelve partially assembled SNAREpins which were clamped in this state by either one (Syt1 or Syt7) or two synaptotagmins (Syt1/Syt1 or Syt1/Syt7). The dynamics of each Synaptotagmin C2 domain were described by the Markov kinetic schemes shown in Figure 4B using the parameters we previously constrained. We assumed that *k_on_* was limited by diffusion to 1 µM^-1^ ms^-1^ and *k_off_* = 150 ms^-1^ based on the intrinsic Ca^2+^ affinity *K_d_* = 150 μM, which is similar for both Syt1 and Syt7 C2 domains^20,45,46^. *k_in_* = 100 ms^-1^ based on the characteristic time for Synaptotagmin C2 domain rotation and membrane insertion^47^. *k_out_* = 0.67 ms^-1^ for Syt1 and *k_out_* = 0.02 ms^-1^ for Syt7, determined from the apparent rates of C2 domain dissociation from lipid membranes measured in the presence of EGTA using stopped-flow experiments^46,48,49^. In reaction Scheme 1 (Figure 4B), we considered that C2 domain membrane insertion leads to the instantaneous release of the fusion clamp. In reaction Scheme 2 (Figure 4B), we assumed that a delay between the Ca^2+^-triggered membrane insertion of the Syt7 C2A domain and the removal of the fusion clamp is described by a first-order reaction with rate the rate *k_u_* = 0.0002 ms^-1^ (this value was chosen as it best fit the experimental data within the tested *k_u_* range of 0.01 – 0.0001 ms^-1^). We further assumed that the clamp could be restored following membrane dissociation at an identical rate *k_c_* = 0.0002 ms^-1^. The rate of vesicular fusion was determined by assuming that the repulsive forces between a docked vesicle and the plasma membrane amount to an energy barrier E_0_ = 26 k_B_T^50^. Once both of its Synaptotagmins are in an unclamped state a SNAREpin contributes ΔE = 4.5 kBT of work towards overcoming this energy barrier^51^. With *n* uninhibited SNAREpins the fusion barrier is spontaneously overcome through thermal fluctuations at a rate given by the Arrhenius equation 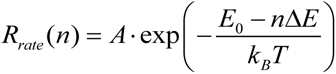, where the pre-factor A = 2.17×10^9^ s^-1^ considering that a single SNARE complex can mediate fusion *in vitro* on a timescale of 1 sec^47,51^. Monte Carlo estimates of the cumulative probability of vesicle fusion in response to a Ca^2+^ activation signal were derived from at least 1000 stochastic simulations of individual vesicles in all scenarios. We estimate that this restricts prediction error due to stochastic variation to less than 1%. All simulations were carried out in Python 3.10. To accommodate the observed variability in the timing of Ca^2+^ signal arrival, which can span up to three frames under our experimental conditions (see Supplementary Figure 4), we applied a temporal blurring of the model output, by smoothing the data across a time window of 0.45 sec, equivalent to the duration of three imaging frames.

