## Supplementary Figures for "A minimal presynaptic protein machinery mediating synchronous and asynchronous exocytosis and short-term plasticity"

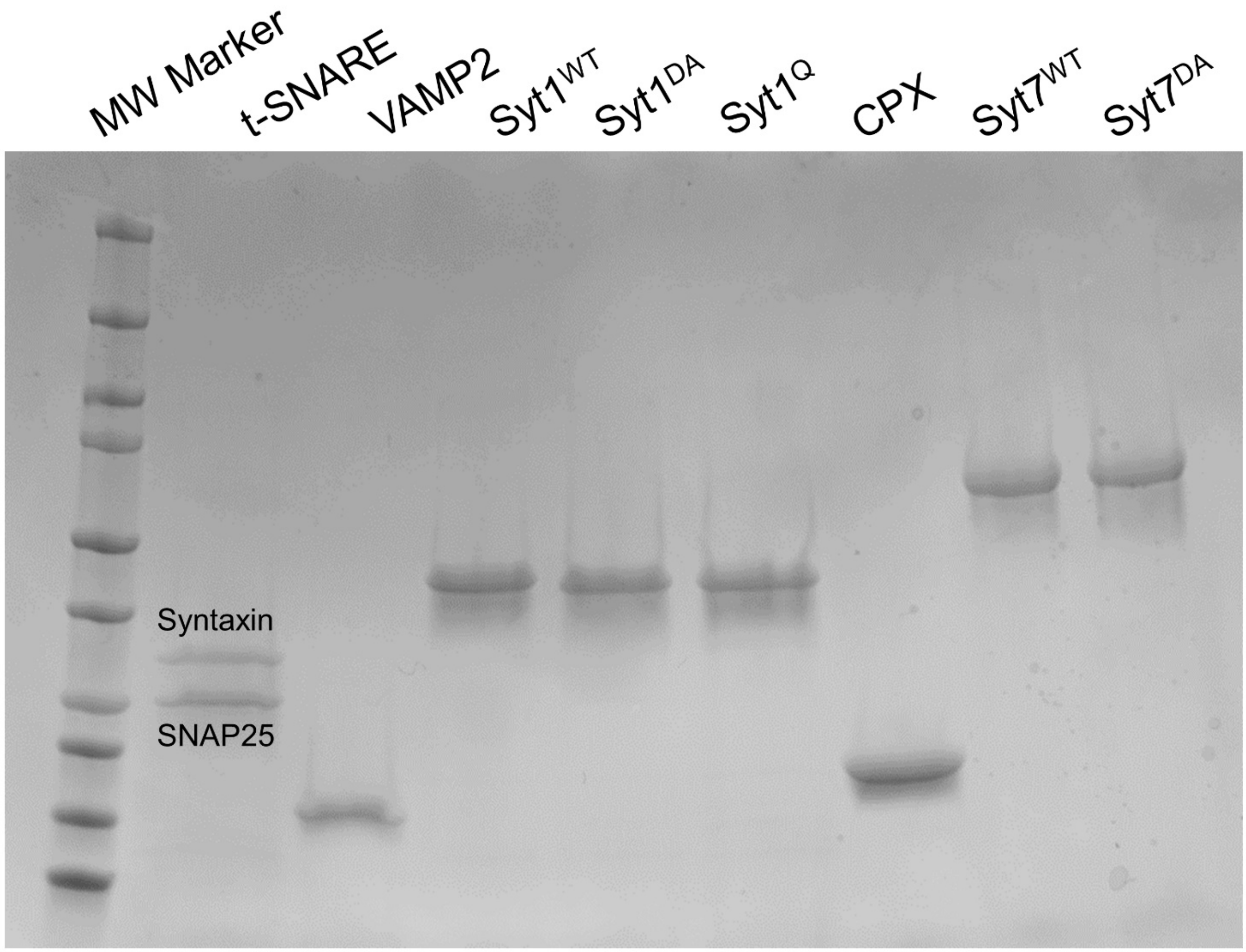


**Supplementary Figure 1.** Coomassie stained SDS-PAGE gel image of the proteins used in the report. All proteins were expressed and purified using the bacterial expression system as described in the Methods section.

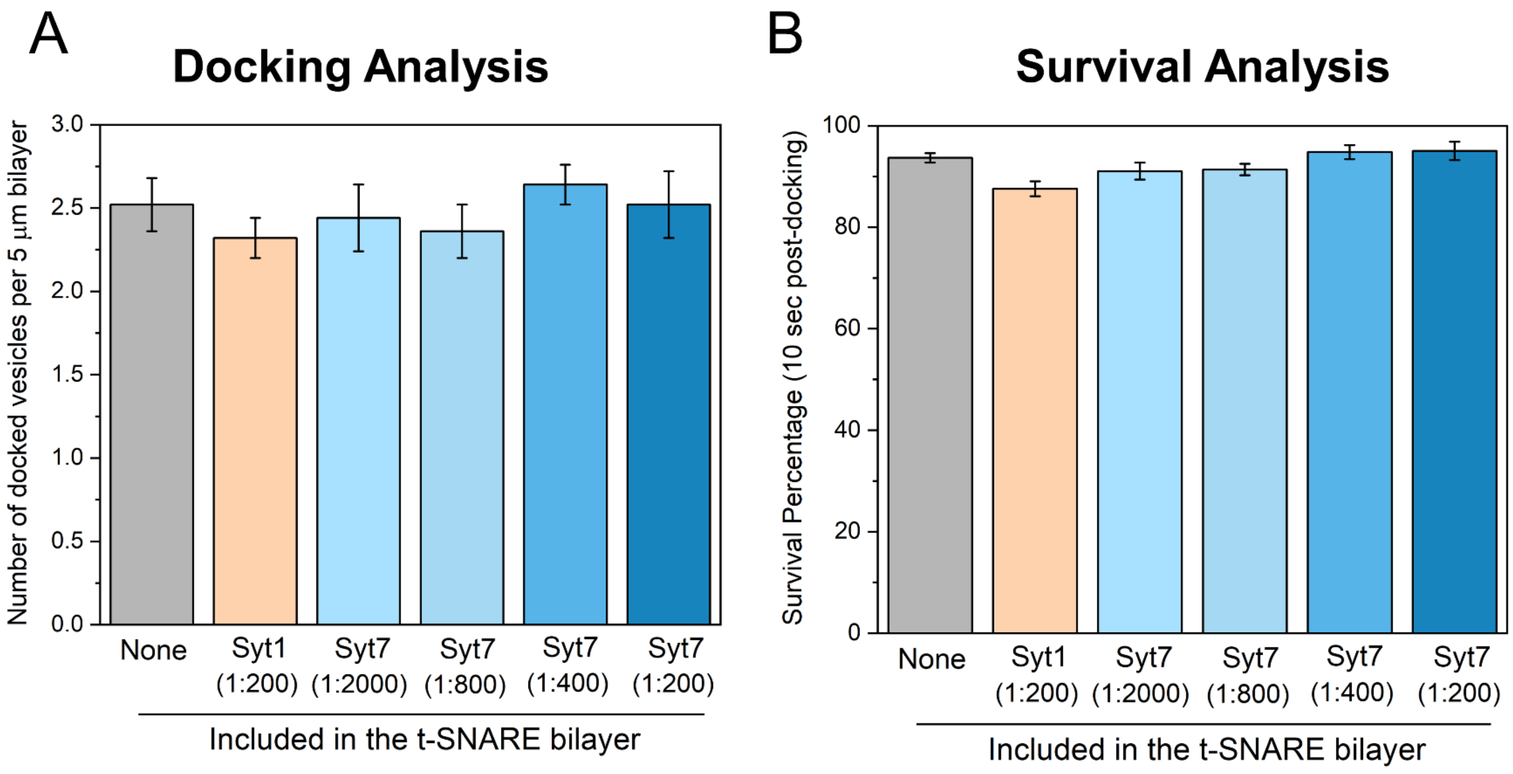


**Supplementary Figure 2.** Effect of Syt7 on the docking and fate of the docked vesicles under resting conditions (A) The inclusion of Syt7 in the bilayer did not affect the docking of the Syt1/VAMP2 vesicles, with a similar number of vesicles stably attaching to the bilayer under all conditions. (B) The majority (>90%) of the docked vesicle, irrespective of Syt7 presence, remained immobile and un-fused *i.e.*, stably 'clamped' at resting conditions. Data (mean ± standard deviation) are from 5 independent experiments for each condition (~ 40-50 vesicles per experiment).

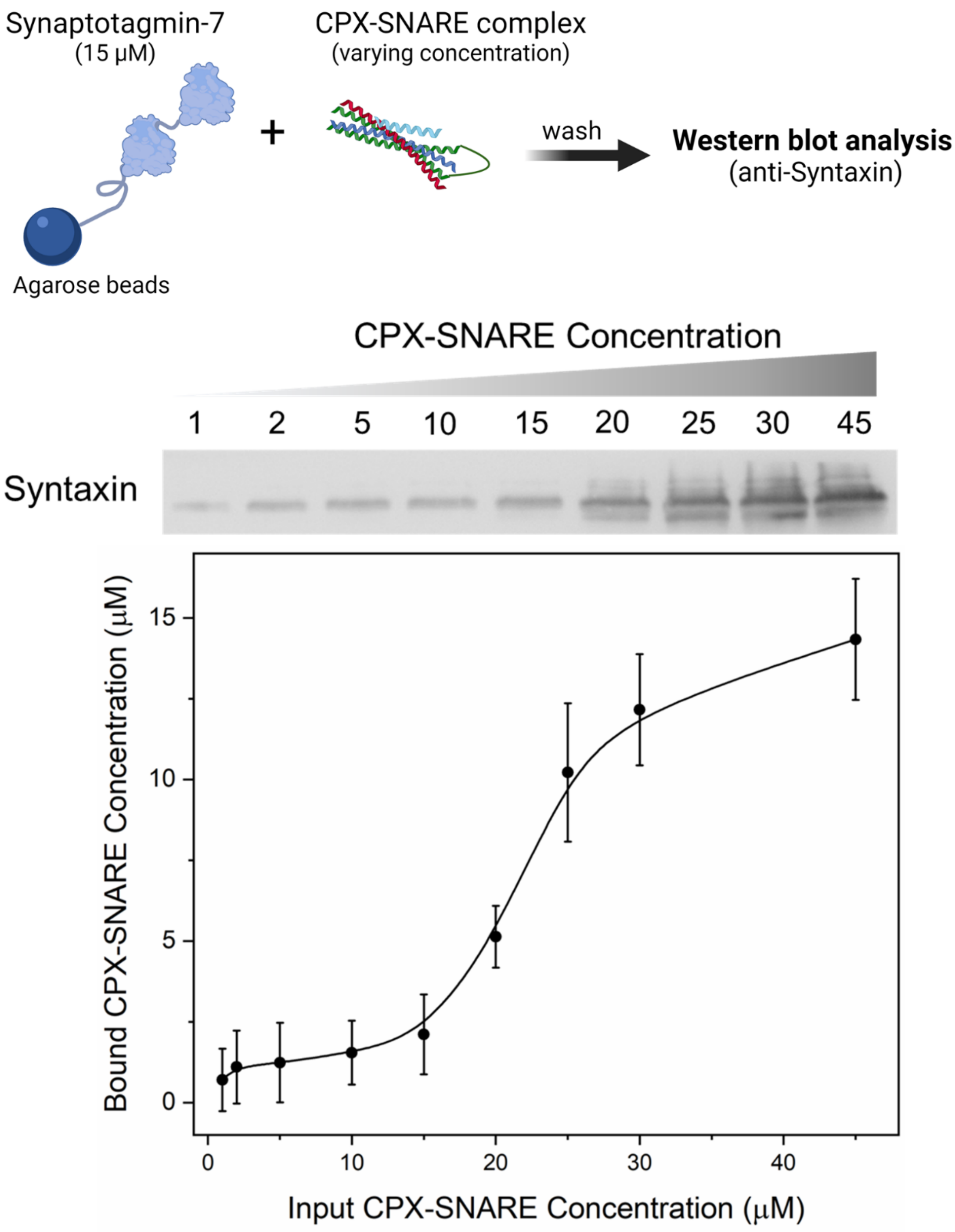


**Supplementary Figure 3.** Direct interaction between Syt7 and SNARE complex was examined using the pull-down assay with Syt7 as 'bait' and CPX-SNARE complex as 'prey'. The amount of SNARE bound was assessed using quantitative western-blot analysis with anti-Syntaxin antibody. We detected an increasing amount of CPX-SNARE bound to Syt7-containing agarose beads, with saturation observed around 30 μM input concentrations. Data from 3 independent pull-down experiments are shown.

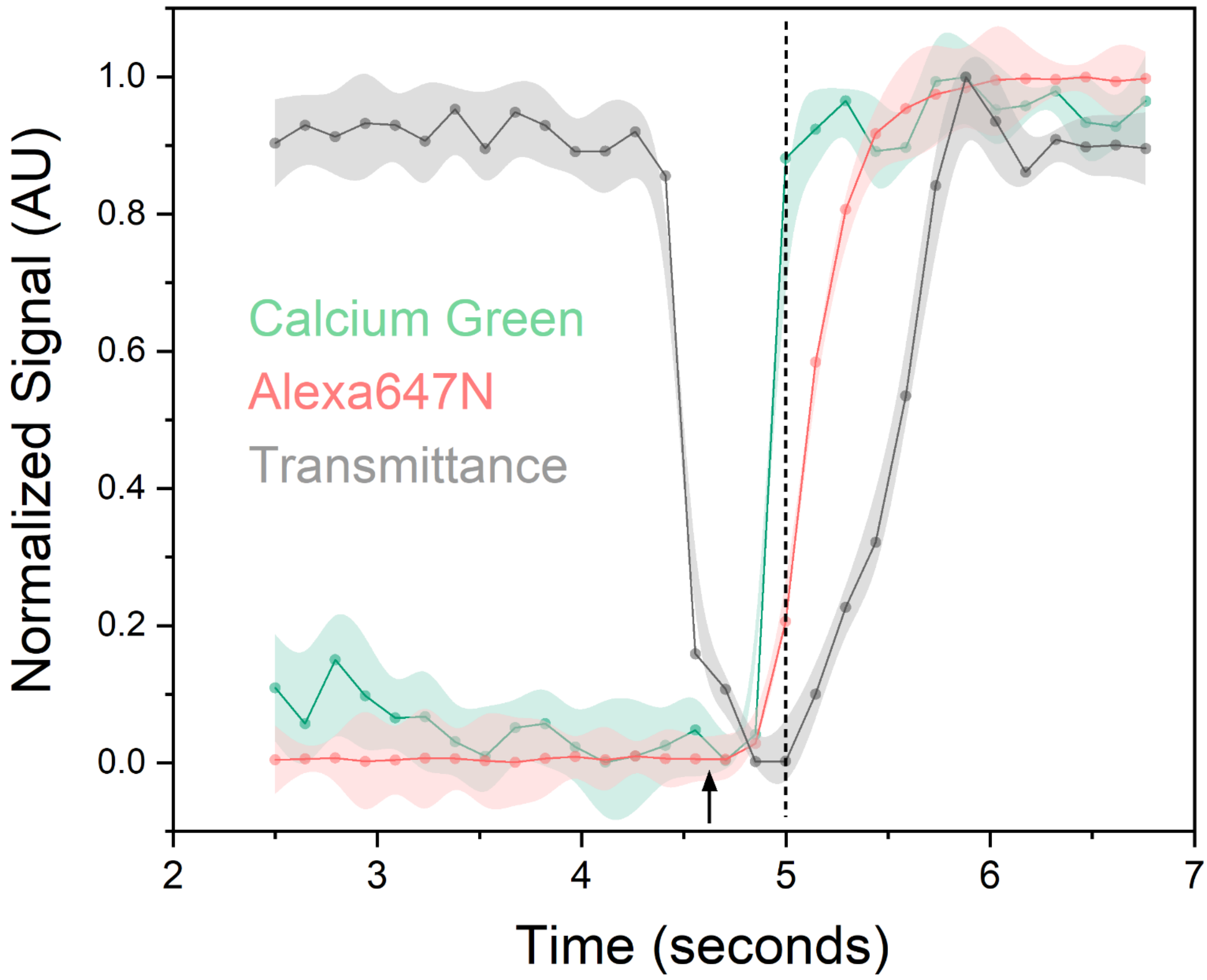


**Supplementary Figure 4.** To examine the $Ca^{2+}$-evoked fusion of docked vesicles, we added $CaCl_2$ (final [$Ca^{2+}$] of 100 µM in the experimental buffer) into the chamber, as denoted by the arrow. In all experiments, we used the transmittance signal to estimate the arrival of $Ca^{2+}$ at the bilayer as it correlated with the minima of the transmittance signal (grey curve). For select cases, we also included a high-affinity $Ca^{2+}$ sensor, Calcium Green C24 (green curve), in the bilayer to track the arrival of $Ca^{2+}$ at the bilayer directly. We confirmed that it corresponds to the changes in the transmittance signal. Since Calcium Green fluorescence saturated within a single frame, in the calibration experiments we used the soluble Alexa647 dye mixed with 100 µM $CaCl_2$ as a 'tracer' to track the actual diffusion of $Ca^{2+}$ into the chamber (red curve) and thus estimate [$Ca^{2+}$] kinetics at docked vesicles.

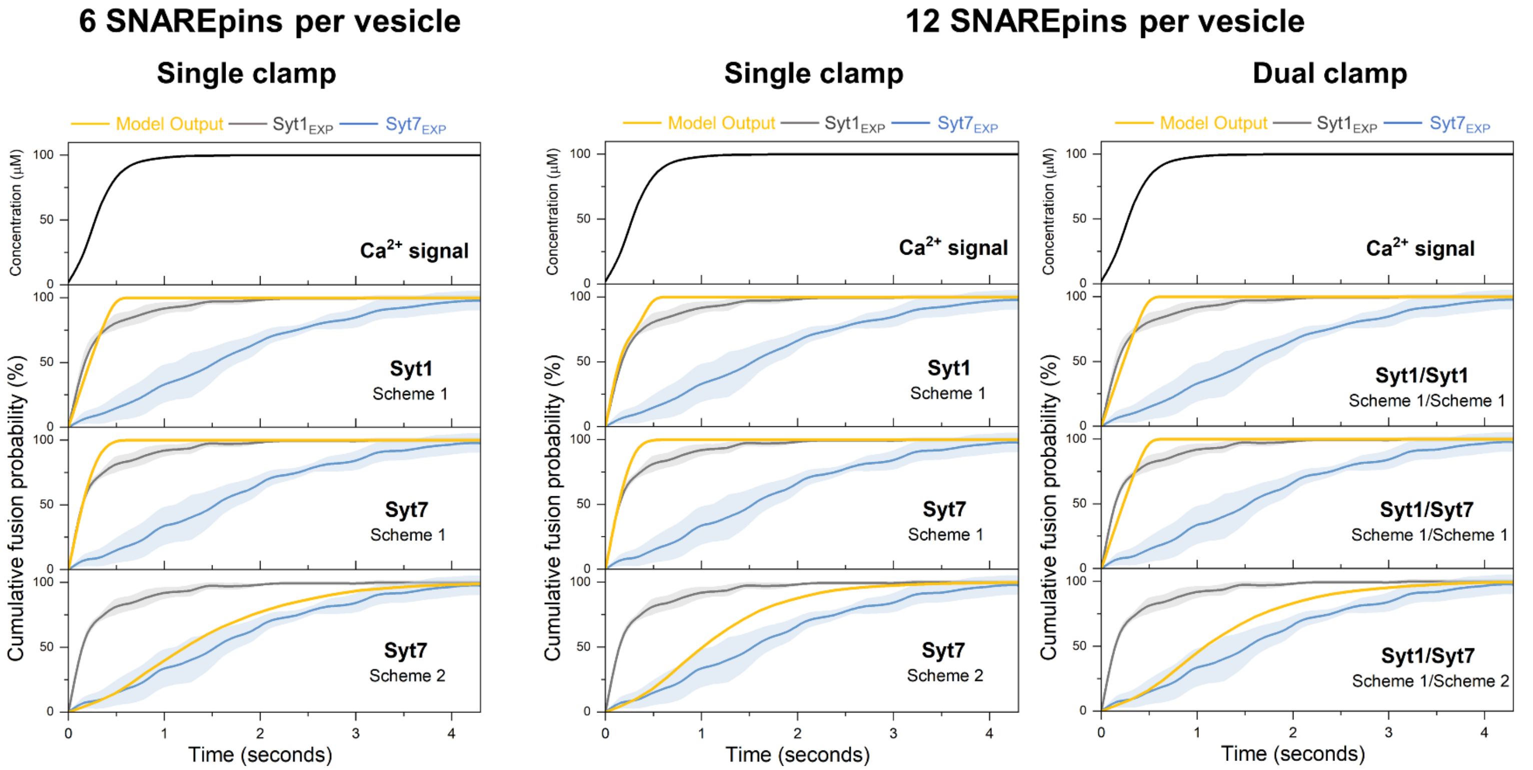


**Supplementary Figure 5.** The time course of vesicular fusion (Model Output) was simulated in response to the experimentally constrained $Ca^{2+}$ signal (Supplementary Figure 4) for models with different numbers of SNAREpins and clamp architecture. As illustrated in Figure 4, we tested different kinetics of clamp reversal with Scheme 1 assuming instantaneous removal of the Synaptotagmin fusion clamp upon membrane insertion, and Scheme 2 assuming a delay. Experimental data (from Figure 1) for the $Ca^{2+}$-triggered fusion of Syt1 containing vesicles in the absence ($Syt1_{EXP}$) or the presence of saturating levels of Syt7 ($Syt7_{EXP}$) are plotted for comparison. The model suggests that observed fusion kinetics can be explained by the mechanism with differential rates of fusion clamp removal for Syt1 (instantaneous) and Syt7 (delayed), regardless of the number of SNAREpins per vesicle or the single vs. dual synaptotagmin fusion clamp architecture. For each modeled condition a minimum of 1000 stochastic simulations were performed to compute the average response.

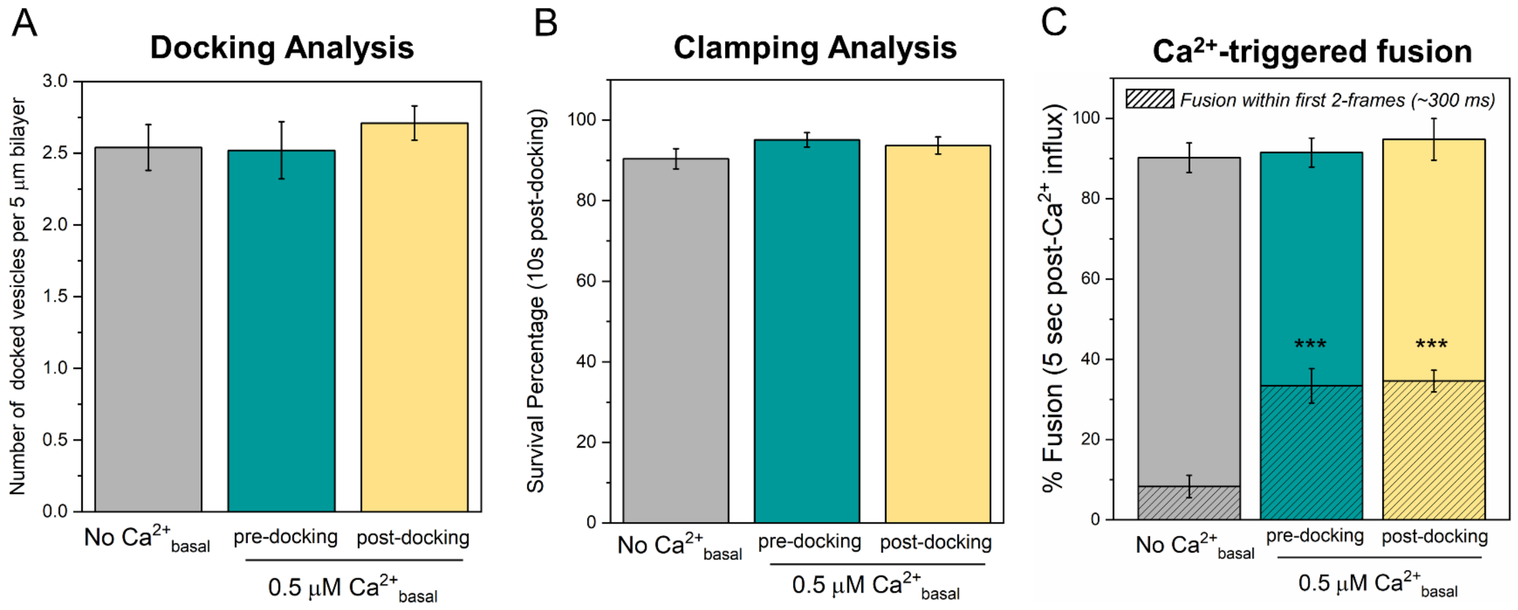


**Supplementary Figure 6.** The effect of elevated $[Ca^{2+}]_{basal}$ and the order-of-addition on vesicle docking, clamping, and $Ca^{2+}$-evoked fusion under $Syt1^{WT}/Syt7^{WT}$ conditions. (A, B) Increasing the $[Ca^{2+}]_{basal}$ to 0.5 µM during (green) or after (yellow) docking of the vesicles had no effect on number of vesicles docking or the fate of the docked vesicles as compared to the control (grey) condition with zero $[Ca^{2+}]_{basal}$. Similar number of docked vesicles were observed, and the majority remained stably clamped. (C) Elevating the $[Ca^{2+}]_{basal}$ enhanced the $Ca^{2+}$-evoked synchronous fusion component (hatch portion), with no observed difference based on order-of-addition of 0.5 µM $[Ca^{2+}]_{basal}$. This data demonstrates that Syt7 facilitates synchronous fusion of docked vesicles. All experiments were conducted using Syt1/VAMP2 vesicles, with Syt7 (1:200) included in the t-SNARE containing suspended bilayer. Data (mean ± standard deviation) are from 3 independent experiments for each condition (~ 40 – 50 vesicles per experiment). *** $p<0.001$ using the student's t-test.

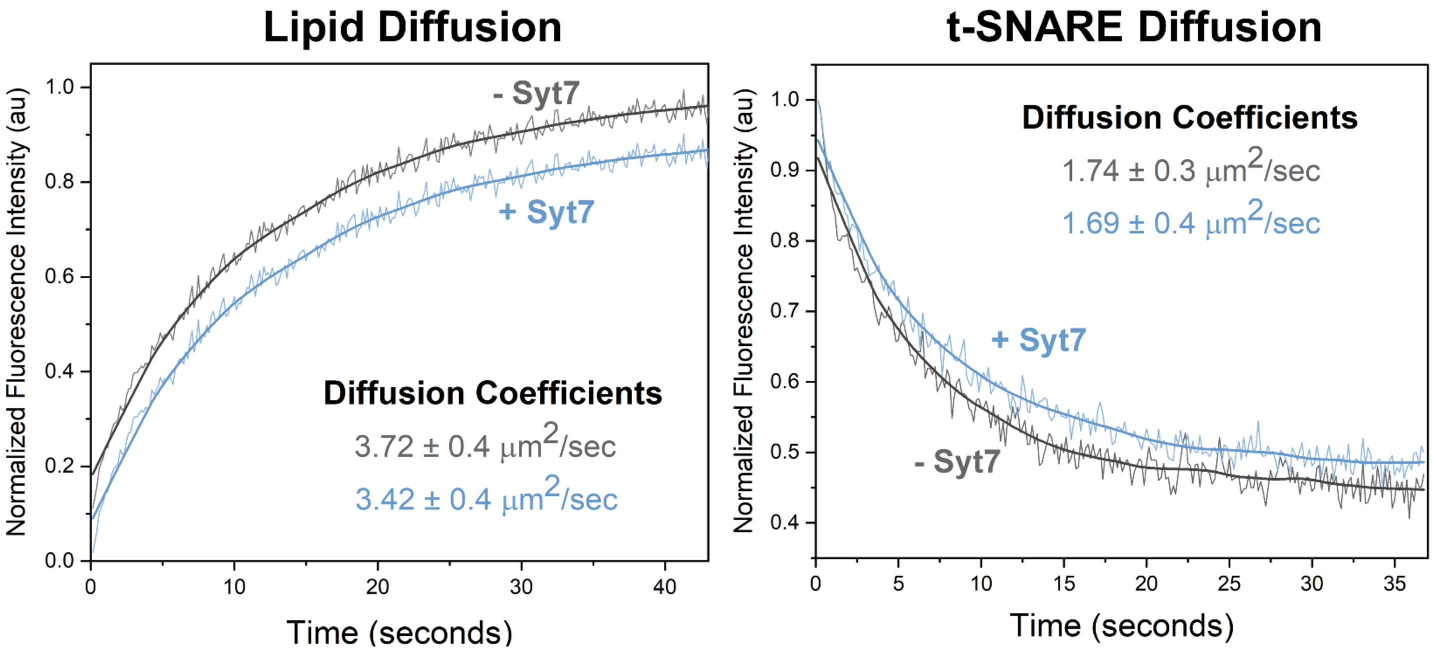


**Supplementary Figure 7**. The diffusion of lipid and t-SNARE on the suspended lipid bilayer is not affected by the inclusion of Syt7. The fluidity of the suspended lipid bilayer containing t-SNARE (Syntaxin/SNAP25) without or with Syt7 (1:200 protein-to-lipid ratio) was measured using fluorescence recovery after bleaching (FRAP) analysis and was analyzed on Wolfram Mathematica using a custom script to estimate the diffusion coefficient as described previously[1,2]. We used the fluorescence signal from ATTO465-PE (2%) included in the bilayer for lipid diffusion and Alexa568-labeled t-SNAREs reconstituted on the bilayer to track the protein diffusion.

### *Supplementary References*


1 Ramakrishnan, S., Bera, M., Coleman, J., Rothman, J. E. & Krishnakumar, S. S. Synergistic roles of Synaptotagmin-1 and complexin in calcium-regulated neuronal exocytosis. *Elife* **9**, doi:10.7554/eLife.54506 (2020).

2 Ramakrishnan, S. *et al.* High-Throughput Monitoring of Single Vesicle Fusion Using Freestanding Membranes and Automated Analysis. *Langmuir* **34**, 5849-5859, doi:10.1021/acs.langmuir.8b00116 (2018).
